# Macroevolutionary dynamics of climatic niche space

**DOI:** 10.1101/2021.12.09.471977

**Authors:** Ignacio Quintero, Marc A. Suchard, Walter Jetz

## Abstract

How and why lineages evolve along niche space as they diversify and adapt to different environments is fundamental to evolution. Progress has been hampered by the difficulties of linking a comprehensive empirical characterization of species niches with flexible evolutionary models that describe their evolution. Consequently, the relative influence of external episodic and biotic factors remains poorly understood. Here we characterize species’ two-dimensional temperature and precipitation niche space occupied (i.e., species niche envelope) as complex geometries and assess their evolution across a large vertebrate radiation (all Aves) using a model that captures heterogeneous evolutionary rates on time-calibrated phylogenies. We find that extant birds coevolved from warm, mesic climatic niches into colder and drier environments and responded to the K-Pg boundary with a dramatic increase in disparity. Contrary to expectations of subsiding rates of niche evolution as lineages diversify, our results show that overall rates have increased steadily, with some lineages experiencing exceptionally high evolutionary rates, associated with colonization of novel niche spaces, and others showing niche stasis. Both competition- and environmental change-driven niche evolution transpire and result in highly heterogeneous rates near the present. Our findings share the limitations of all work based purely on extant taxa but highlight the growing ecological and conservation insights arising from the model-based integration of increasingly comprehensive and robust environmental and phylogenetic information.

## Introduction

Unravelling the evolutionary tempo and mode that over millions of years have brought about the substantial variation in niche space occupied by species worldwide is a foundational goal in biology [1, 2]. Climatic niches, the multidimensional hypervolume of climatic conditions in which species can persist [3, 4], represent a key component of this niche space. While recent years have seen important progress in the establishment of phylogenetic frameworks and the characterization of species climatic niches, we still lack a comprehensive understanding of the macroevolutionary dynamics of climatic niche evolution.

Niche divergence, whereby species adapt to a different set of environmental conditions than their ancestors, is prevalent in nature. While under a neutral expectation lineages would diverge from one another in niche space by random drift, with niche disparity across lineages increasing linearly with time, several constraints and fundamental mechanisms predict a contrasting pattern. For instance, if the total niche space available is finite, then a natural upper bound to dissimilarity among species is expected to be met eventually. Biological constraints, such as those imposed by physiology, mutation, or generation time might slow or prevent the spread towards certain regions of niche space [5]. “Red Queen Hypothesis” [6] type explanations emphasize biotic drivers of evolutionary change [7], with competitive dynamics resulting in either spatial exclusion or, among coexisting lineages, an initial acceleration of niche evolution driving niche partitioning [8]. In finite geographical and ecological space this recurrent spatial and niche partitioning is expected to increasingly hinder niche evolutionary rates as clades diversify and suffer from density dependence [1, 9]. In the absence of such constraints, however, competition among an increasing number of lineages could instead accelerate niche evolutionary rates [10]. Niche shifts might be associated with diversification in other ways. For instance, punctuated equilibrium predicts that bursts of evolutionary change occur at speciation [11]. Considering that most speciation events in terrestrial groups such as tetrapods are allopatric, incipient species will inhabit environments that differ to varying degrees, thus stimulating adaptation [12].

Another set of “Court Jester” type hypotheses argue for environmental changes as the prevalent evolutionary force [7]. For example, environmental fluctuations are expected to speed up evolutionary rates as species adapt to new ecological opportunities [13]. Extreme events such as the Cretaceous-Palaeogene (K-Pg) transition, triggered ubiquitous diversification and caused the preferential extinction or survival of certain phenotypes, acting as ecological filters for surviving lineages [14, 15]. And it brought about a sudden worldwide winter and a devastation of forest habitat, with few lineages surviving on spatially scattered refugia and exposed to freezing temperatures and dry weather [16]. The resulting massive extinction, dispersal and population fragmentation, accompanied by selection for small body sizes, enhanced speciation and adaptive potential rates on the lineages most tolerant to colder, drier conditions that survived and diversified [14, 17].

## Results

Here we use a novel two-dimensional niche framework for phylogenetic analysis to characterize the joint macroevolutionary dynamics of thermal and precipitation niche domains across all extant lineages of birds. In contrast to previous analyses, our approach is the first to assess niche evolution using the full polygon occupied by each species across the two-dimensional temperature and precipitation space [4], which, for clarity, we henceforth call the species climatic niche ‘envelope’. Figure 1b illustrates this difference: for each species we inform the model with their occupied niche envelope rather than simply using single niche coordinates (say, the average value across each axis). A species niche envelope results from the union among all composing individuals, whereby each individual’s envelope is expected to occupy a substantial portion of the species envelope [18]. Scaling-up, each species’ climatic envelope occupies a substantial portion of available climatic niche space, with high degree of overlap with other species’ envelopes, manifesting the need to take into account the whole of the polygonal surface.

**Figure 1:**
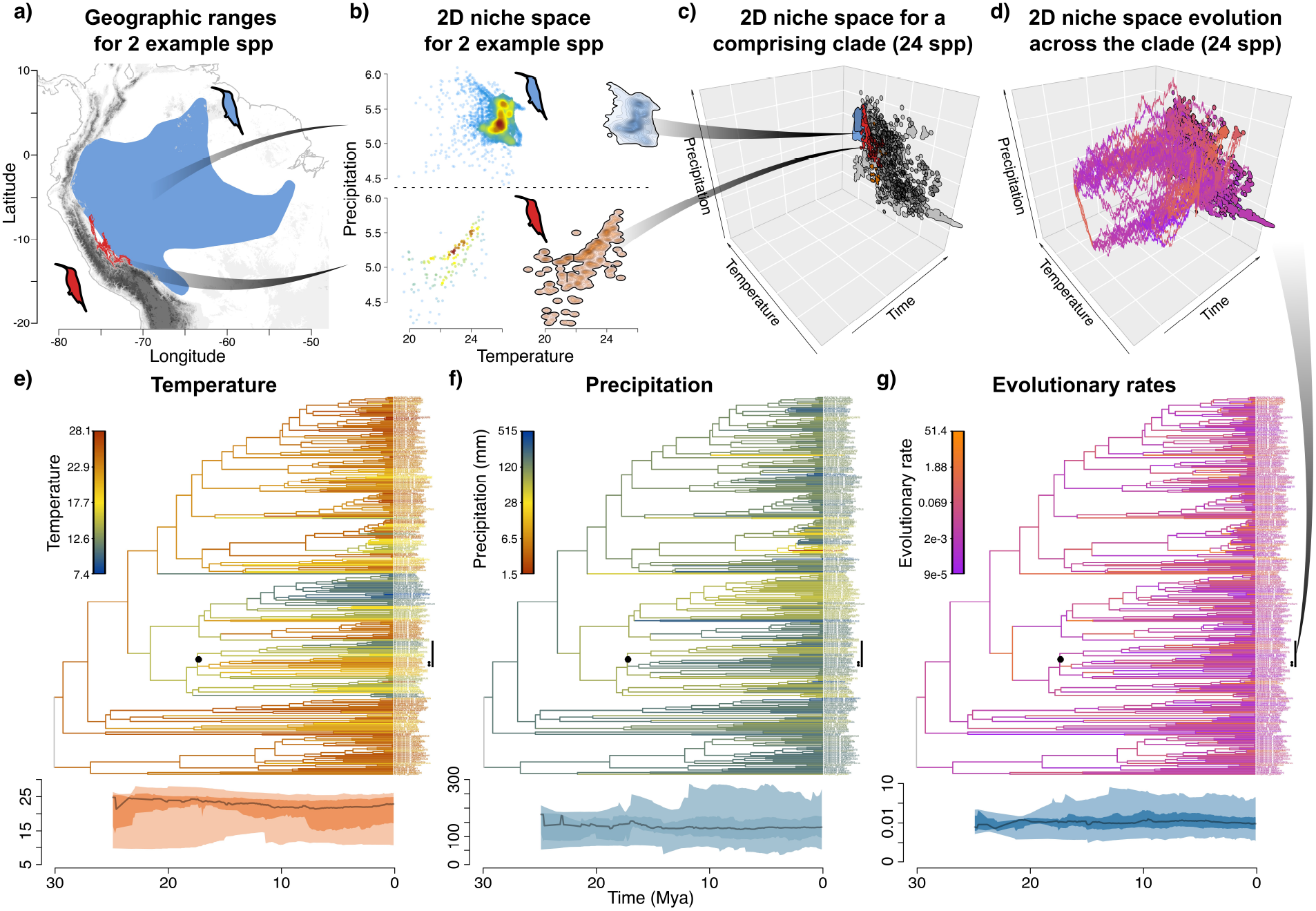
Niche reconstruction approach illustrated for hummingbirds (Trochilidae) **a)-d)** Demonstrate for two species (Heliodoxa aurescens, blue; Heliodoxa branickii, red) the steps from spatial data to climatic niches and their reconstruction. a) Shows their geographic ranges, corrected for elevation preferences. b) Characterizes the density of temperature and precipitation conditions experienced over each range grid cell. Left shows raw densities (with lighter colors indicating lower density) and right the corresponding 95% Gaussian kernel density estimates (with shading and contour lines corresponding to each 0.05 density level). c) Climatic niches for all 24 species of a comprising clade mapped as the present-day states across time and d) with five posterior samples resulting from the niche ancestral estimation (branch colors represent rates as in g). **e)-g)** Estimated climatic niches and their evolutionary rates (expected squared difference in traits change over time, described by the Brownian motion diffusion rate matrix Σ) for all hummingbirds (Trochilidae). Shown are a single MCC phylogeny (top) and across all its branches over time (bottom, line corresponds to median, and darker and lighter shading to 50% and 95% quantiles, respectively. Graphs provide the ancestral estimates from the niche space evolution model for e) posterior median temperature, f) posterior median precipitation, and g) posterior median branch-specific evolutionary rates. The highlighted subclade in c) & d) is marked with a black vertical line and a circle around the shared ancestral node. 32

Considering climatic niches as envelopes thus prevents the loss of the highly complex multivariate and within-species variation of climatic niches caused by their reduction to coordinate-point estimates (e.g., [19, 20]), which compromises ancestral and rate estimates of niche evolution [21]. Specifically, we extracted for each species their realized temperature and precipitation during the breeding season and used Gaussian kernel density estimation to delineate their occupied climate niche space as polygonal envelopes in bivariate climate space (Fig. 1a-c). We then modelled evolution of species climatic niche by performing inference over polygonal envelopes rather than point coordinates, making use of the ‘Relaxed Random Walk’ model, a flexible and relaxed model of multivariate trait evolution that allows each branch in the phylogeny to have a specific niche evolutionary rate [22]. This model allows the detection of variable evolutionary rates across branches, clades and time, without an *a priori* expectation of the temporal or taxonomic behaviour of evolutionary rates (Fig. 1d,g) [22], thus avoiding a smoothing temporal and taxonomic heterogeneity typical for most models of trait evolution, but see other approaches allowing variable rates [23]. For clarity, we consistently use the term evolutionary rate to refer to the expected squared difference in traits change over time (*i*.*e*., ^◦^C for temperature and mm/month for precipitation), which corresponds to the multivariate Brownian motion diffusion rate matrix Σ (see Methods). For the example of hummingbirds (Trochilidae; Fig. 1e,f), a clade inhabiting a gamut of environments, this phylogenetic climate estimation uncovers that their broad range of occupied conditions extends to ancestral lineages, and that clade-level aggregates remain relatively constant over time.

To comprehensively describe the multivariate niche evolutionary history we estimate cross-branch temperature and precipitation trends (Fig. 2a-d) and two-dimensional niche occupancy of all ancestral lineages over time (Fig. 2f & Supplementary Video 1). Across all birds, and only reflecting lineages with extant descendants, our model estimates an ancestral origin of birds in warm, mesic environments with limited disparity across clades throughout the Cretaceous (Fig. 2ab,f, Supplementary Fig. 3bc). While the biogeographic origin of modern birds (Neornithes) is hypothesized to be southern (West Gondwana), where our reconstructed environment was prevalent, biogeographic evidence remains ambiguous as to the location or climate inhabited by the Most Recent Common Ancestor (MRCA) of modern birds [24]. The climate inferred at the origin of birds and subsequent climate evolution agree with the, compared to today, more modest latitudinal temperature gradient further back in bird evolutionary history and, possibly, higher extinction rates in colder regions [25, 26].

**Figure 2:**
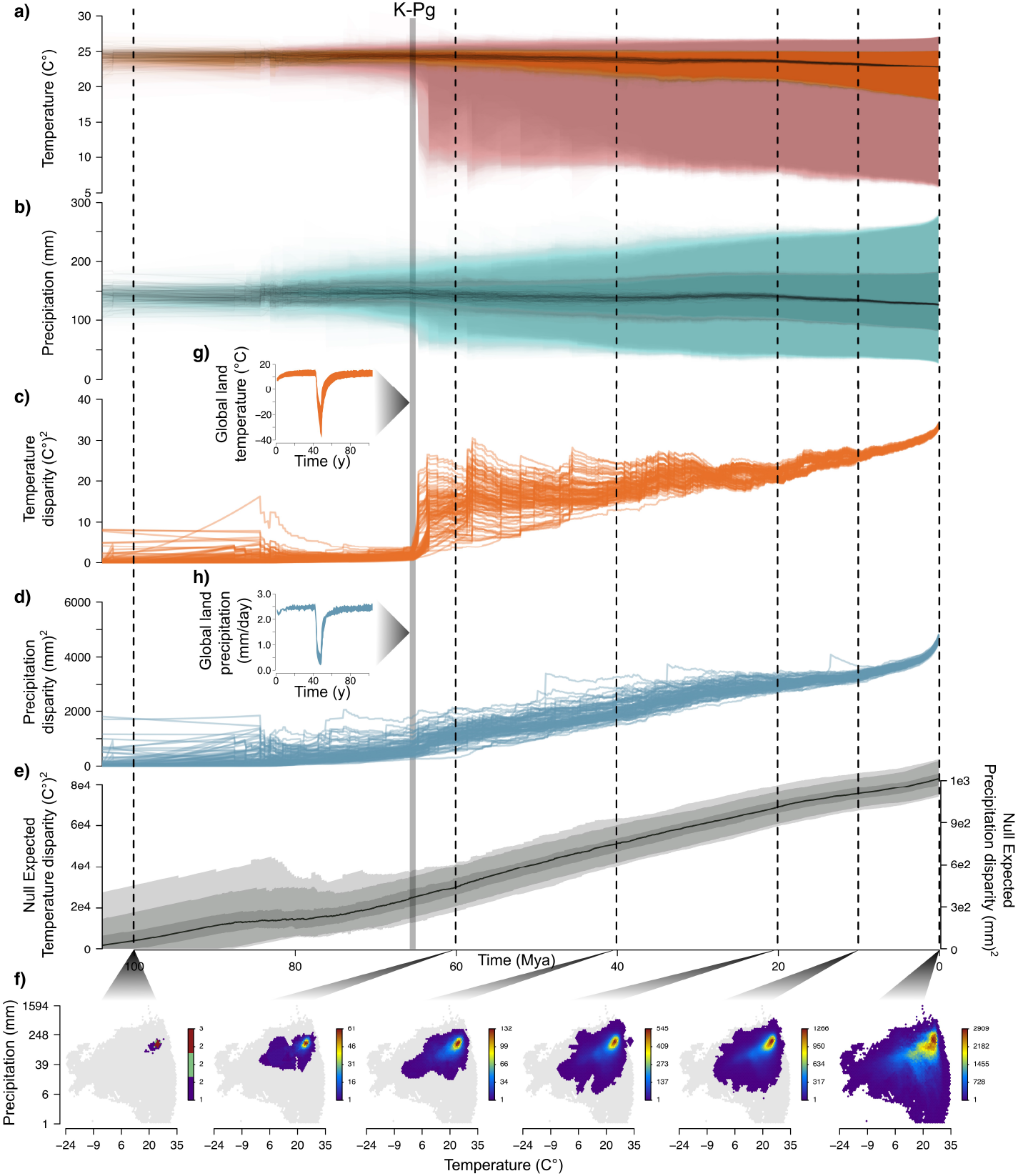
Evolution and disparity in bivariate climatic niche space in extant birds. **a)-d)** show reconstructed temperature and precipitation values and disparity across coexisting branches over time for 100 tree samples. Middle line corresponds to the median and darker and lighter shading to 25 − 75% and 2.5 − 97.5% quantiles of branch values, respectively. **e)** Expected disparity given a model of strict Brownian motion and tree topology across the full set of 100 empirical birds trees used in this study. **f)** Time slices at 100, 60, 40, 20, 10 and 0 Mya of environmental space showing niche filling. The hexagonal grid is drawn by summarizing each lineage’s climatic niche as a 95% highest posterior density (HPD) polygon and coloring each grid cell by with the count of overlapping climatic polygons at a given time slice. As a point of reference, the currently occupied realized climatic space for all birds is shown in grey background. **g) & h)** Range of simulated scenarios of the effect of the K-Pg asteroid winter impact on global land temperatures and precipitation (following ref [16]).

Coinciding with the aftermath of the Cretaceous-Paleogene (K-Pg) mass extinction, at around 66.02 Mya, the range of predicted ancestral conditions and their disparity, measured as sample variance in ancestral climatic estimates across contemporary lineages, dramatically increased, with lineages colonizing colder and modestly drier environments (Fig. 2a,b). This shift is strongly visible in the temporal patterns of niche disparity, measured as sample variance in ancestral climatic estimates across contemporary lineages. We calculated null expectations for niche disparity given the number of species and tree topology during this time under a diffusion process of trait evolution (Fig. 2e) and under models of bounded evolution (Supplementary Fig. 7), which reflect the limited range of climatic conditions available at a given time on earth. We find that the sudden increase after the K-Pg boundary that deviates substantially from these expectations, and especially so for temperature. We note that this result emerges only from the signal carried by the evolutionary history of niche evolution across the tree, since our model does not explicitly account for episodic events of disparity. This suggests that the observed divergence is not the result of new lineages, but rather of specific colonizations of unoccupied niche space. The concurrence of this climatic niche expansion with the K-Pg Boundary is consistent with the “ big-bang” model of diversification wherein extinction followed by the emergence of new environments and habitats gave rise to new ecological opportunities in surviving lineages [14, 17]. Indeed, recent tip-dating efforts on subsections of the bird tree did show an increase in speciation rates after the K-Pg [17], and it is concordant with observations of posterior dispersal towards the cooler northern hemisphere [27].

After this rapid and substantial niche expansion around the K-Pg, we find climatic disparity increasing in near linear fashion, with a final recent sharp increase (*<* 4 Mya; Fig. 2c,d), which is not expected by the number of lineages alone. We hypothesize that this acceleration towards the present is due to the partition of a shared fundamental niche among young radiations that suffer from competitive exclusion following allopatric speciation. Recently diverged species are expected to retain common environmental tolerances while occupying a subset of their climatic niche given the geographic partition of their ancestor, potentially leading to over-differentiated realized climatic niches [28]. Similarly, observed niche differences at these shorter time scales (*<* 4 Mya) might represent differences only in their realized rather than fundamental niche space, which would not necessarily involve divergent adaptation. Evolutionary changes that result from changes in the realized niche can only be detected over such short time scales and are expected to suffer from the observed higher evolutionary rates than those changes involving the fundamental niche (Fig. 3a). This difference between the realized and fundamental niche is expected to decrease as lineages adapt to the new conditions. Alternatively, as past environmental fluctuations shaped available climatic space [29], lineages might have been responding evolutionarily to the emergence of novel climates [30]. Indeed, temperate and boreal climates are relatively new with their area expanding towards the present [31]. Finally, while boundary effects alone can create a final sharp increase in disparity (Supplementary Fig. 7), the empirical increase is much shallower than that expected under the null model, suggesting that other, intrinsic biological factors are at play.

**Figure 3:**
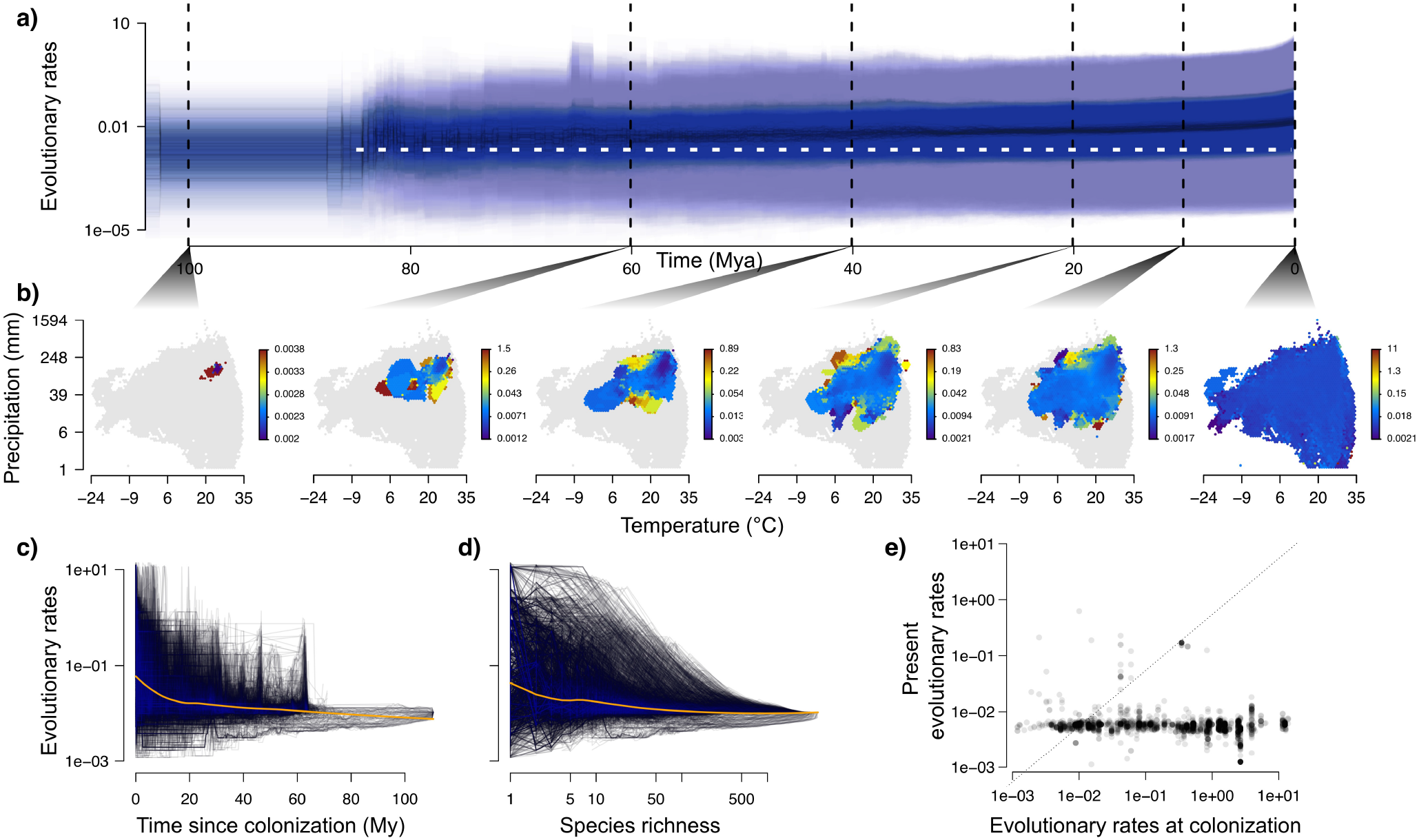
Rate of climatic niche evolution over time. **a)** Estimated branch-specific evolutionary rates across coexisting branches along time. The horizontal white dashed lined marks rate constancy, in contrast to the empirical pattern (other graph details as in Fig 2). **b)** Time slices showing niche filling with hexagonal grid cells colored by the median of the evolutionary rates among the overlapping lineages at a given time slice. **c)-e)** show median evolutionary rates for lineages intersecting with a hexagonal grid of niche space (see b). c) relates median lineage rates in each niche-space cell to the time since first colonization of that cell (i.e. first switch from ‘grey’, background, to ‘color’ in b)). d) relates for each niche-space cell the changing median rate to the changing count of lineages in that same cell. e) Compares the median initial rates in each niche-space cell to its present rate (line shows 1:1 relationship).

We find that the continuous increase in disparity is paralleled by a general increase of climatic evolutionary rates through time (Fig. 3a). Rates, however, vary enormously across branches and consistently so through time (Fig. 3, 4 & Supplementary Fig. 3a), implying that at any given time, there are lineages experiencing exceptional high rates and others undergoing very low evolutionary rates. Branch scalars for median rates of bivariate evolution range from about 9 × 10^−5^ to 51 in hummingbirds (Fig. 1g) and from 2 × 10^−6^ to 500 in all birds (Supplementary Fig. 3a). This seems to conflict with expectations derived from early-burst theory, where high initial rates of trait evolution are followed by stasis as niche space gets progressively occupied [1, 9]. However, by simultaneously visualizing evolutionary rates and how niche space has been progressively occupied (Fig. 3b & Supplementary Video 2), we find that lineages experience high rates of bivariate niche evolution as they move into new niche areas, with rates subsequently waning in descendant lineages not experiencing substantial niche shifts (higher rates are assigned to larger niche shifts, Supplementary Fig. 6b,c).

**Figure 4:**
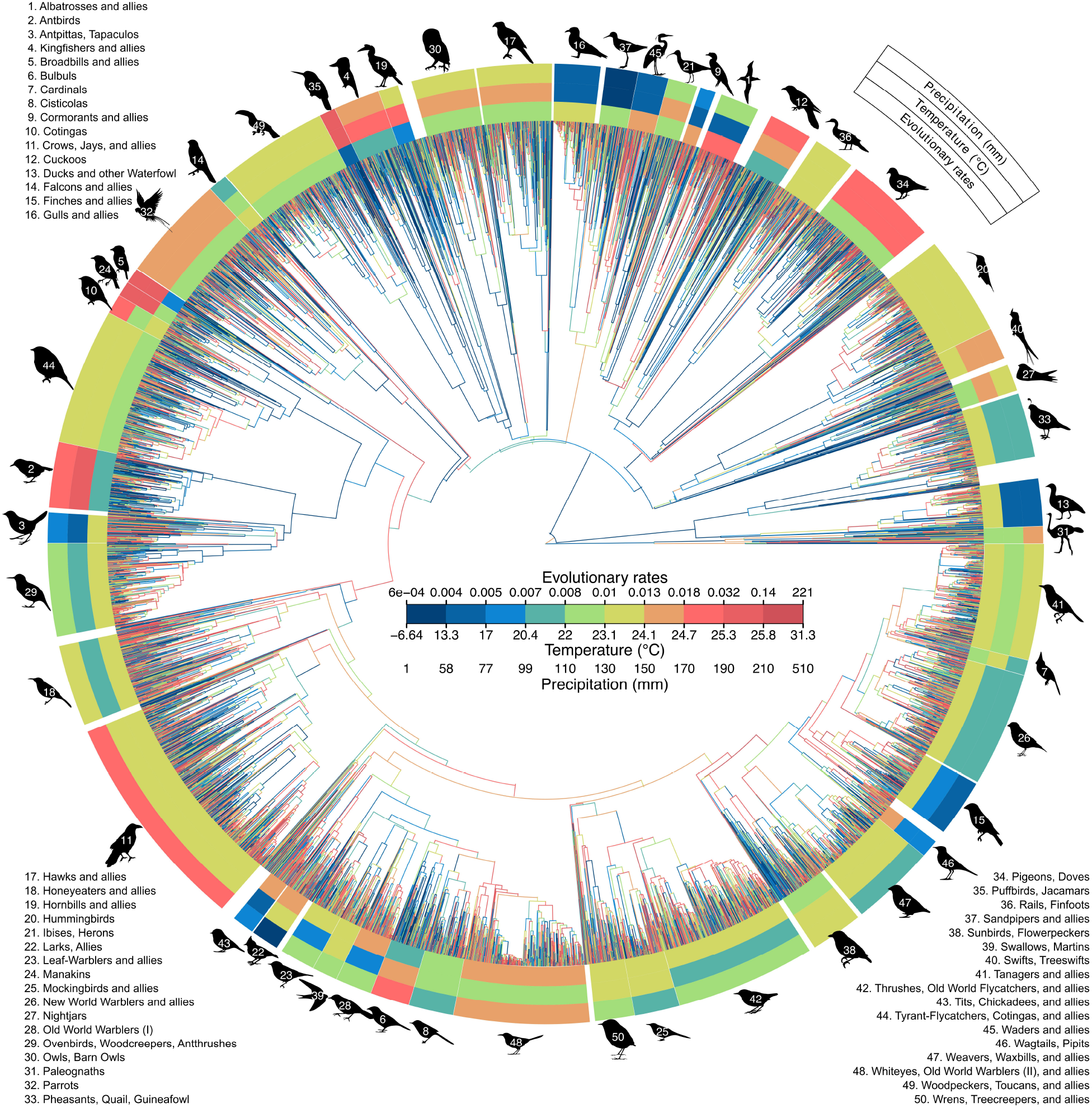
Variation in reconstructed bivariate temperature and precipitation niche space and associated evolutionary rates across extant birds. Shown is a Maximum Clade Credibility tree with branches colored according to median branch evolutionary rate scalar (*ϕ*_*b*_). For visual clarity, each color represents a quantile and do not represent equal magnitude. Outer circles represent clade medians for, from inside to outside, evolutionary rates, temperature and precipitation. Silhouettes characterize major clades (holding at least 50 species).

Finally, we partition temperature and precipitation niche space into discrete (hexagonal) units to assess how rate dynamics vary across niche space over time. We find that after new niche space is colonized and followed by a concomitant increase in the number of lineages, there is a strong slowdown of evolutionary rates (Fig. 3c,d). Consequently, present rates are lower than initial rates in ca. 84% of cases (Fig. 3e). Lastly, we find that evolution across bivariate climatic space is not independent, but rather that adaptation to warmer temperatures is associated with humid environments and vice-versa (posterior median correlation = 0.51, 95% highest posterior density interval = [0.48, 0.54], Supplementary Fig. 3a).

## Discussion

These results highlight the importance of phylogenetic scale in assessing niche evolution. At small phylogenetic scales, our results are consistent with Simpson’s [1] model wherein species experience high evolutionary rates as they enter new adaptive zones but become progressively slower as niche space saturates [32]. When scaling-up to broader phylogenetic scales, contemporary subclades undergo recurrent evolutionary phases of niche evolution as they diffuse through climatic niche space, resulting in both high and low evolutionary rates at any moment in time (Fig. 3 & 4). Across the full avian tree of life, an intriguing picture of replicate filling of environmental niche space emerges, with select lineages retaining their ancestors’ niches and others occupying particular sections of temperature and precipitation space over short time scales. While larger clades generally show similar niches over time, reflective of conservatism, the underlying evolutionary rates exhibit strong heterogeneity (Fig. 4).

Our results suggest that, while populating niche spaces decreases subsequent evolutionary rates at short timescales, sufficient environmental opportunity has been available for young radiations to colonize new (or recycle old) adaptive zones, thus maintaining (and increasing) a set of high rates across time. These results are remarkably consistent with simulated scenarios where inter-specific competition increasingly accelerates niche evolutionary rates as lineages diversify in an effectively boundless environmental space [10]. Likewise, our results are in line with previous analyses of morphological evolution across time which have not detected any slowdown as lineage richness increases [23, 33]. Considering the bounded nature of Earth’s temperature and precipitation and the observation that rates of climatic evolution increase together with temperature and precipitation disparity, either more time is needed for this radiation to face density effects, or most clades are not moving into new climatic spaces but recolonizing already occupied environmental space. Alternatively, an increased availability of new environmental space could explain accelerated evolutionary rates as lineages adapt to previously unavailable climates [31]. Different realized climatic niches result only from dissimilar spatial distributions, yet the latter does not assure the former. The seemingly relative ease at which species can overlap across climatic niche space could be explained by spatial segregation as well as differentiation along other ecological and behavioral axes facilitating coexistence at finer scales [34].

The evolutionary rates presented in this paper encompass a significant larger range, by orders of magnitude, than those previously reported in the literature (Fig. 4, SI). Very fast niche shifts are commonly reported over ecological time scales (but see [35]), but rarely emerge from fossil and phylogenetic analyses at evolutionary timescales [28, 36]. Notably, a methodological dependence exists when a single evolutionary rate for a tree is used (smoothing underlying rate variation) and ancestor-descendant change is divided by the corresponding branch length: as the interval at which change is measured increases, estimated rates tend to decrease, particularly in methods using strict Brownian motion or similar models where rates of change are proportional to branch length [19, 37]. Our relaxed random walk model overcomes these methodological constraints by explicitly untangling branch lengths from evolutionary rates (explaining < 0.01 of the variation in climatic rates, slope = −0.21, SE = 0.012, Supplementary Fig. 6a). By explicitly considering bivariate climatic space, our analyses also minimize biases that arise from collapsing the multidimensional and multi-population nature of species climatic niches to single points, as typical for earlier rate estimates [21]. Most of our recent evolutionary rate estimates are, on average, higher than those previously suggested (median 95% temperature rates range = 3.43 − 1177.9 ^◦^C^2^ My^−1^), yet, even the maximum median branch rates remain below projected changes of climate change (SI). Interpreting these past estimates of evolutionary adaptation to novel climates as signal for future adaptability to projected rates of temperature and precipitation change thus presents an overall adverse picture. Nonetheless, responses will be highly species and habitat specific, and, coupled with the large rate heterogeneity observed, might offer hope and highlight the potential for some lineages much more than others to cope with impending change.

Despite the methodological advances, our results, like all work based purely on extant taxa, remain limited by their ignorance about extinct species and paleoclimatic information, which are not represented in our reconstruction of climatic niche evolution. Comprehensive fossil information could prevent potential biases in our climatic estimates if extinction probability is associated with species’ climatic niches, and it would provide germane information about past rate dynamics and ancestral states [38]. For instance, what would seem like available niche space using extant species-only reconstructions could actually have been occupied by now extinct clades [33], most significantly so across the K-Pg Boundary where substantial diversity was lost after the mass extinction [39]. This issue also affects the interpretation of our estimated evolutionary rates since inclusion of extinct lineages would subdivide a branch (and its associated rate) and, since the total amount of niche divergence to explain can only be equal or larger, evolutionary rates should either increase or balance a decrease in one branch with an increase in another. Similarly, like all multivariate models of trait evolution, we assume that available climatic space is limitless (with precipitation being non-negative), yet available climatic conditions during birds’ evolutionary history have had bounds that fluctuated substantially [29]. Non-absorbing bounds, as expected from evolution across climatic niche space, pose an upper bound to inferred evolutionary rates because, after some time, the amount evolutionary change measured cannot be larger than the breadth of available niche space [19]. Similarly, species evolving near the boundary of climatic conditions could have their rates of adaptation not be as fast as they could be given unbounded space. Indeed, the steady increase in evolutionary rates towards the present could also respond to surge of available cold, dry climatic conditions lineages could adapt to. As sampling issues of the fossil record ameliorate, its inclusion can dramatically improve inference [38]. We expect that sufficient fossil evidence in the future combined with improved models of paleoclimatic conditions [16, 29] will provide an important test, and update, of our phylogenetic work on extant species and will at that point further advance of our understanding of biological climatic niche dynamics.

Our analyses used comprehensively characterized bivariate climatic niches, phylogenetic information with uncertainty propagation and a flexible model of evolution to identify dramatic among-branch variation and consistent increase in evolutionary rates throughout the radiation of birds. Our results suggest an evolutionary model of recurrent events where lineages move into new niche spaces at high evolutionary rates followed by relative stasis as species diversify within. This process seems to have continued at an increasing rate even when niche space is bounded, suggesting that species can recolonize already occupied expanses of niche space at a steady pace, at least for the duration of the Aves radiation. Why some lineages maintain and conserve their ecological niche while others quickly adapt to new optima across environmental space might be due to different abiotic or biotic factors, including rare extreme events (such as the K-Pg boundary), physiology, coexistence dynamics, and dispersal opportunities. Usually regarded as antipodes, our results support an interplay between the Red Queen, by agreeing with expectations from ubiquitous competition-driven niche evolution, and Court Jester hypotheses, by the significant effect of environmental catastrophes. We find that complex niche dynamics reiterate consistently throughout the evolutionary history of birds, but overall, estimated rates are low compared to what might be required to adapt to impending future changes. With ever increasing rates of climatic change forecast for the planet, further research expanding a multi-dimensional understanding of niche evolution to other groups and phylogenetic scales is urgently needed. Increasingly comprehensive and robust environmental and phylogenetic data and models offer new opportunities for understanding the evolutionary dynamics at macroevolutionary scales, informing biogeography, ecology, and conservation.

## Methods

### Estimating bivariate climatic niche envelopes

#### Climatic niche envelope as a polygonal domain

Here we follow Hutchinson in considering a species niche as an *n*-dimensional volume or envelope, Ξ^*n*^, mapped onto environmental space [4] and focus on two of the most ecologically relevant abiotic variables, temperature, *T*, and precipitation, *P*\. We consider an environmental Cartesian plane *℘* with yearly average temperature as the *x*-axis measured in ^◦^C (plainly referred as temperature for simplicity), and average monthly precipitation as the *y*-axis measured in mm/month in logarithmic scale (plainly referred as precipitation for simplicity). Precipitation is a monthly basis rather than annual because, as detailed below, for migrant species we only considered breeding months. One can measure these climatic variables across a species distribution range and map them on environmental plane *℘*. Let **C**_(*m*,2)_ be a matrix representing *m* climatic coordinates for a given species, such that

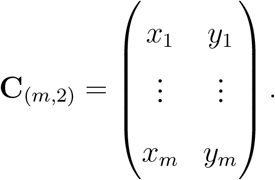

These *m* coordinates represent samples from the present-day species’ realized niche, a subset of the intersection of their fundamental niche and available environmental space [30].

Characterizing the fundamental climatic niche space is challenging, particularly for all species within a large radiation (i.e., involving experimental manipulation), yet we contend that modelling the evolution of the realized climatic niche is still pertinent (or even more so than the fundamental niche). Since the realized niche is a non-biased subset of the fundamental niche (i.e., the arrangement differences between the realized and fundamental niches have an expectation of 0), it should be a non-biased representation of climatic niche space evolution. Or, put simply, there is no expectation that the realized niche is a subset that tends to be, say, towards the higher values available in the fundamental niche for any niche axis. Expectedly, interacting closely-related species partition a shared fundamental niche, resulting in non-biased species-specific realized niches, as remarkably evinced in elevational replacements across mountains of young radiations [40, 41]. Secondly, species adapt to conditions within their realized niches [42]. Thus, climatic niche evolution results from the microevolutionary selective pressures on the realized niche which ultimately dictate the macroevolutionary dynamics of the fundamental niche [42]. The degree in which an observed niche shift is due to explorations within the fundamental niche or a shift in the latter will be inversely related to time. Thus, we expect to infer higher evolutionary rates at shorter time scales (i.e., towards the present) where shifts in the realized niche have still not prompted the evolution of the fundamental niche but acknowledge our inability to distinguish between these.

Our goal is to define the two-dimensional niche envelope along temperature and precipitation, 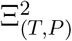, such that it describes the current environmental conditions where species can persist and proliferate [43]. We approximate 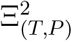 by delineating a two-dimensional envelope **D** in *℘* composed of one or more non self-intersecting polygons, which may contain holes [44]. This envelope should have the following desired property: if we define the intervals *I*_*x*_ = max{*x*} − min{*x*} + *ϵ*_*x*_, where {*x* : *x* ∈ **C**_(*,1)_} and *ϵ*_*x*_ is measurement uncertainty in *x*, and, similarly, *I*_*y*_ = max{*y*} − min{*y*} + *ϵ*_*y*_, where {*y* : *y* ∈ **C**_(*,2)_} and *ϵ*_*y*_ is measurement uncertainty in *y*, then Area(**D**) ≤ *I*_*x*_*I*_*y*_. That is, we would like to not only independently describe the optimal range along the axes, but their interaction [43].

To construct **D**, we assume that the probability that an individual from a given species occurs within *℘* is not uniform, but instead some areas concentrate higher likelihood of occurrence than others [30, 45]. This reflects on the concept that fitness for a given species varies within *℘* and our goal is to find the envelope **D** wherein species have 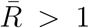, where 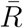 is the *per capita* growth rate [42, 43]. Given that we are restricted to the realized niche, we follow [30] in considering that 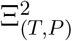 results from a species response-surface of density across different populations. Thus, we assume that **C**_(*m*,2)_ constitute *m* samples drawn from a common unknown density function *f*_Ξ_(*x, y*), which corresponds to a fitness gradient wherein 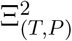 delineates the *T* and *P* values where 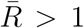. To estimate *f*_Ξ_(*x, y*), we use bivariate kernel density estimation with a Gaussian kernel as implemented in the *kde2d()* function of the MASS package for R [46, 47]. Kernel bandwidth selection will affect the resulting density, and we discuss our selection below. We then delineated **D** by considering the 95% highest density region of our estimated density, 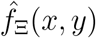 (i.e., there is a 0.95 chance that a point drawn from 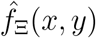 would fall within **D**). That is, **D** specifies the (*x, y*) values such that 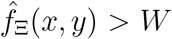 where *W* satisfies 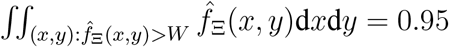.

Understanding and delineating the niche 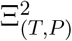 using **D** has several advantages over other methodologies. Specifically, we are able to obtain polygonal surfaces of high complexity that characterize the particular distribution of all niche coordinates along *℘*. In this way, convoluted boundaries and holes within a species niche space are easily characterized [44, 48] and we do not assume a uniform distribution across the niche coordinates during the estimation (such as when using, for instance, a convex hull), but we take into account the density of points along *℘* to define **D**. This not only agrees with the notion of fitness landscapes along environmental space [43], but also helps in avoiding bias by disregarding possible inaccuracies (e.g., large deviants) in our niche coordinates.

#### Avian realized climatic data

We extracted **C**_(*m*,2)_ for all 9, 993 bird species, following the taxonomy of [49], using the following procedure. First, we obtained expert distribution range maps for each species. These range maps have been validated at a resolution of *ca* 110 km [50], describing species climatic distributions at a relatively low resolution if one considers, in particular, strong climatic gradients, such as in mountainous regions. To increase the accuracy of our climatic niche envelopes, we therefore incorporated elevation distributions, thereby significantly enhancing the resolution of presence-absence information in expert distribution ranges [51]. We used a recently compiled dataset that includes data for species that inhabit in mountainous regions (ca. 86%) and, whenever available, uses mountain region specific elevational information, for full description see [51]. The remaining species do not inhabit strong elevational gradients and thus their niche accuracy should not be compromised by it. A global Digital Elevation Model (DEM) was obtained at a resolution of 10 minutes latitude/longitude from [52]. We then refined the expert distribution ranges by removing all cells that had an elevation value outside from where the species occurs.

Climatic data was extracted at a resolution of 10 minutes longitude/latitude from [52]. The use of a ten-fold higher resolution for climatic data compared to that at which planar range maps have been validated (i.e, 1^◦^longitude/latitude) is justified by the incorporation of threedimensional presence-absence information [51]. For each grid cell within the refined species breeding distributions we extracted the average annual temperature and average monthly precipitation. For migratory species, we only considered the temperature and precipitation during the breeding months when extracting climatic information (i.e., April-October for Northern Migrants and October-April for Austral Migrants). 230 species (*ca*. 2.3% of the total) were left without climatic information. Of these, 144 species inhabit small islands where climatic information was not available in [52] and we thus used the climatic characterization of islands with area *>* 1 km^2^ from [53] instead.

The remaining 86 species (*ca*. 0.86% of the total) are narrow ranged species for which the elevation refinement procedure described above reduced their range completely. The cause was the resolution of the DEM used, which represents a summary average over deviant values within grid cells at a 10 minute grain. Thus, we used the 30 arc-seconds resolution NASA/NGA Shuttle Radar Topography Mission DEM to refine again these species according to their elevational range. The use of a higher resolution layer for these species is justified by its necessity: our information on where these species occur is at a higher resolution than that of our previous layers after incorporating the species’ elevational ranges. To obtain average annual temperature and monthly precipitation information at this resolution we used data from CHELSA (Climatologies at high resolution for the earth’s land surface areas) [54]. Using higher resolution (30 arc-seconds) for these 0.86% of species with narrow elevational ranges (compared to the 10 minutes latitude/longitude resolution from [52] used for most species), helps avoid averaging over the steep environmental gradients that they inhabit.

#### Estimation of avian bivariate climatic envelopes

Our final climatic dataset for each bird species is a collection of *m* grid cells with bivariate climate coordinates that make up a matrix **C**_(*m*,2)_. Given that our climatic evolution model (see below) assumes that the reconstructed traits do not have bounds, we added one unit to the precipitation data and transformed by taking the natural logarithm. This is only done to precipitation data since in contrast to temperature data, it cannot be negative. We mapped **C**_(*m*,2)_ into *℘* and used bivariate kernel density estimation (see Fig 1b,c). We assumed a common bandwidth across all species so that each niche coordinate has the same weight, independent on the range size of the species (and thus of the resulting number of climatic coordinates *m*). Optimal bandwidth selection is usually performed by minimizing the discrepancy between the estimator 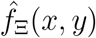 and the true density *f*_Ξ_(*x, y*) [55]. We attempted to use optimizers that minimize a measure of discrepancy such as the Mean Integrated Square Error (MISE), however, the large amount of data points (*>* 6.8 × 10^7^ environmental coordinates) made the computation impractical. Thus we first visually inspected the use of different kernel bandwidths, and decided to use a 1% of the total axis range across all species striking a good balance between boundary delimitation and complexity for species with large *m* and cohesion for species with small *m*. Across all species, the annual temperature range is [−23.55, 34.7] in ^◦^C and the average monthly precipitation range is [0.0, 7.33] in ln(mm/month), resulting in a bandwidth of *ca*. 0.58 and *ca*. 0.07, respectively. We then explored two commonly used approximations to an optimal kernel bandwidth when assuming that *f*_Ξ_(*x, y*) is a multivariate normal, Table 4.1 in [55]. The resulting bandwidths where similar to those estimated by visual inspection (Silverman’s rule: *ca*. 1.5 and *ca*. 0.13 for temperature and precipitation, respectively; Scott’s rule: *ca*. 0.47 and *ca*. 0.04 for temperature and precipitation, respectively). Since the resulting polygons are comparable, but the 1% bandwidth estimates lie in between the other approximations, we only used the envelopes **D** that resulted from using the visually inspected bandwidth. To gauge the consequences of this decision, we evaluated the impact of using different bandwidths on evolutionary rates (see below).

### Trait evolution for niche spaces

To model the evolution of climatic niche envelopes, we used the bivariate Relaxed Random Walk (RRW) described in [22, 56] along a phylogenetic tree. This model relaxes a time-homogeneous Brownian diffusion process by multiplying tree-wide evolutionary rates with a scalar *ϕ*_*b*_ for each branch *b*. Specifically, let Σ be the infinitesimal tree-wide covariance matrix for bivariate diffusion, then the bivariate rate for branch *b* is rescaled to Σ*ϕ*_*b*_, such that the process can vary across branches [22]. We denote 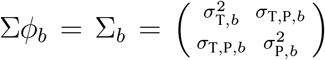. For the model to be identifiable, we assume that *ϕ*_*b*_ follow a Log-Normal prior distribution with mean of 1 and estimated variance *σ*_B_ [22]:

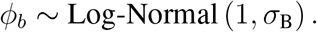

To improve mixing and computational efficiency, the internal nodes, including the root, are analytically integrated for likelihood calculations using multivariate continuous trait peeling as described in [56].

Methods that allow to specify polygonal envelopes instead of points as trait values at the tips have recently been developed [21, 57]. While both of these methods numerically integrate over polygons, [21] integrate a bivariate Gaussian over the polygon to estimate the quasi-exact likelihood and [57] use Markov chain Monte Carlo (MCMC) integration through a uniform transition kernel over the polygonal envelope. Here we numerically integrate over polygonal envelopes following [57], as this then enables us to integrate analytically over all of the internal nodes, improving MCMC efficiency. To further accommodate the size of our dataset, we substantially improved the computational efficiency with which BEAST evaluate the model posterior density if proposed trait values fall within the polygonal domains. We accomplished 8 − 10-fold total run-time improvement through a two-step density evaluation and then memoization, an approach that most MCMC simulators in phylogenetics, including BEAST, have employed to minimize computationally expensive sequence data likelihood evaluations. However, memoization has remained ineffective for the relatively cheap and few prior density evaluations in traditional phylogenetic problems. At scale with thousands of tips, evaluating if Metropolis-Hasting proposed trait values fall within specific polygonal domains, returning a non-zero density, or fall outside, returning zero density, becomes rate-limiting. The two-step procedure first quickly checks if values fall within the minimal hyper-rectangle that encloses each domain. Only when this condition holds does the procedure then more slowly evaluate if the values fall within the domain itself. Finally, memoization of the last set of values that fall within the domain avoid costly re-computation when proposed states are rejected in generating the Markov chain. These extensions are now available in BEAST v1.10 [58].

## Evolutionary analysis of avian climatic niche envelopes

### Phylogenetic information and analysis set-up

Phylogenetic information was based on [49], currently the only source with all extant bird species included as tip taxa in a cohesive phylogenetic framework that uniquely enables the presented clade-wide niche analyses. To properly propagate the phylogenetic uncertainty inherent in this tree, we randomly sampled 100 phylogenetic trees from the posterior distribution and cycle through this set of posterior trees during the MCMC. Ideally, one would perform joint estimation of the tree and RRW model, but this remains computationally unfeasible. For each species we used their polygonal environmental envelope **D** as tip values for trait evolution. Holes within the polygons were accommodated by using “corridors” of 0 area that delineate only one continuous boundary (e.g., see Fig. 1b). Given that our data consist on the realized climatic niche for all species, ignoring dispersal limitations, biotic interactions and behavioural climatic regulation, we acknowledge that our evolutionary analysis do not reflect the evolution of the fundamental climatic niche.

### MCMC run and convergence diagnostics

We simulated two parallel MCMC chains well-passed draws reaching stationarity, which required over a year of computation time at this extreme scale, and convergence was assessed. One chain was run continuously for 378 days on a 3.4GHz core under a x86_64 Darwin Kernel resulting in *ca* 9.86 × 10^9^ iterations. The second chain was run for 601 days on a 1.6Ghz core under a x86_64 Linux Kernel resulting in *ca* 1.15 × 10^10^ iterations. Unfortunately, during the time of our analysis, it was not yet possible to restart the MCMC after termination, hence our two chains. Interested readers can now employ BEAST’s checkpointing capabilities to restart and move chains between computers as necessary. Using visual inspection, we discarded the first 5 × 10^9^ iterations as burn-in and saved every 2 × 10^5^ sample to reduce auto-correlation (see Supplementary Fig. 2). While visually both chains appear to have converged to the same stationary distribution, we used more objective diagnostics to asses convergence using the package coda [59] for R [47]. First, we used the Geweke diagnostic on each of the resulting MCMC chains, which compares the mean from the first 10% of the samples with that of the last 50% and returns the z-score [60]. The chains show plainly that there are no significant differences between means (z-scores in the chain order presented: −0.034 & 0.06822). Secondly, we used the Gelman and Rubin’s convergence diagnostic,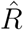, which uses the information from more than one chain by comparing the variance for each chain with that between chains [61, 62]. This diagnostic does not provide a hypothesis test, but, generally, with a value below 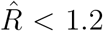 one can confidently accept convergence [63, 64]. For our two chains, 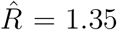 with an upper 95% Confidence Interval = 2.15. We, however, argue that this is a fair diagnostic given our inference problem. Beforehand, one has to consider the data and model size and complexity for the Monte Carlo integration: 19984 branch-specific rates + 19984 ancestral node values + 3 tree-wide evolutionary rates (temperature, precipitation with covariation) + the prior variance, yielding 39972 parameters for each of the 100 posterior trees where tip values are specified using polygons. While we acknowledge that the 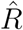 is above the “rule of thumb” threshold, the significant complexity of our highly-dimensional problem and the empirical results that mirror each other from the separate chains strongly suggests that convergence has been reached, and further iterations or chains would likely represent a marginal gain in precision. Similarly, most convergence diagnostic tools are not suitable under limited computational resources with respect to time [65]. Finally, we obtain a minimum of 20 effective sample size (*n*_eff_) across all parameters/chains, which usually are enough to portray a posterior distribution; [64] suggests at least *n*_eff_ *>* 10. Thus we argue that for the results presented here, the macroevolutionary patterns hold between separate MCMC simulation chains and oversuffice in depicting the evolution of climatic space through time for all Aves.

The amount of data and different sources of uncertainty returned by the analysis make elementary summaries of the results challenging without incurring in partial loss of information. For clarity, each posterior sample returns an annotated phylogenetic tree with temperature and precipitation values for each node and a rate scalar for each branch (on top of the tree-wide parameters, Σ, and prior variance). For cross-branch sectional analyses (Fig. 2a,b & 3a-c), we used all the available uncertainty, derived from considering 100 phylogenetic trees in the analyses and the estimates from a final posterior sample of 871 annotated trees. Additionally, we summarized branch-specific results using the 871 annotated trees using ‘TreeAnnotator’ in BEAST, resulting in the maximum clade credibility (MCC) phylogenetic tree with associated posterior distributions for branch-specific scalars *ϕ*_*b*_, which multiply rates of niche evolution, and for ancestral estimates of temperature and precipitation. The product of the tree-wide instantaneous matrix, Σ, with *ϕ*_*b*_ result in branch specific temperature 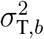, precipitation 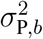, and covariance 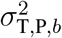 climatic evolutionary rates.

### Sensitivity analysis to phylogenetic uncertainty and bandwidth selection

We explored how phylogenetic uncertainty and bandwidth selection affect parameter inference. We randomly selected a small clade of 18 species that was consistent across 9 pseudo-posterior phylogenetic bird trees and, for each species, estimated **D** using three different bandwidths: 0.1%, 1%, and 2%. For simplicity, we used Range Ancestral State Estimation (“rase”), which assumes strict Brownian Motion while integrating over two-dimensional domains [21], to perform ancestral states estimation for each combination of tree and bandwidth, resulting in a total of 27 different runs. rase makes inference within a Bayesian framework using pseudo-Gibbs MCMC sampling for all derivable parameter conditional distributions combined with Metropolis-Hastings steps to accommodate for arbitrary polygons as tip states [21]. Each analysis was run for 1.5 × 10^5^ iterations, which yielded *n*_eff_ *>* 500 across all parameters for all runs. We note that while we are not integrating over bandwidth selection in our main analysis, we are integrating over phylogenetic uncertainty by using an empirical distribution of 100 trees (see above).

### Comparison to strict and coordinate-only Brownian motion

Finally, we compare our RRW model with a simple multivariate strict BM using the midpoint coordinates of each species climatic envelopes as input. We ran the analyses on the MCC tree and used maximum likelihood for optimization using the ‘mvMORPH’ package in R software [47, 66].

### Expected disparity under Brownian motion without and with bounds

To test if our temporal patterns of disparity, measured as sample variance across time, deviate from a null expectation as given by a strict Brownian Motion model of trait evolution across the set of tree topologies used, we estimated expected disparity through time across the full set of 100 phylogenetic trees. Expected disparity at any time for each tree was estimated following Equation 1 in [67] using the median posterior Σ for the rates. While this expectation is based on strict Brownian motion model, it provides a temporal expectation of disparity under the specific phylogenetic trees used in this study. Deviations from this expectation allows us to conclude that the particular arrangement of tree topologies is not behind the observed patterns. Furthermore, to explore the effect of boundaries in climatic niche evolution, we exployed simulations as an analytical solution to the expected disparity through time was not available. We generated temporal patterns of disparity for temperature and the logarithm of precipitation using 10 simulations for each of the 100 trees using the “phytools” package [68] in R [47]. We used the maximum and minimum current available values of temperature and precipitation on Earth, and used the median posterior Σ for the evolutionary rates.

## Acknowledgments

We thank Hélène Morlon, Daniel J. Field, Michael J. Landis, Julien Clavel, Carlos D. Cadena, Jonathan Drury, Leandro Aristide, Muyang Lu, Shubni Sharma, Richard Li, and, more generally, the Evolvert and Morlon labs for providing feedback on earlier versions of the manuscript. We thank Alfio Alessandro Chiarenza for sharing their data. This project has received funding from the European Union’s Horizon 2020 research and innovation programme under the Marie Skłodowska-Curie grant agreement No 897225 for IQ. This work was partially supported by the National Institutes of Health (R01 AI153044). WJ acknowledges support from NSF DEB-1441737.

## Author contributions

I.Q. and W.J. designed the research and wrote the manuscript. I.Q. and M.A.S developed and conducted the analyses.

## Supplementary Information

Supplementary Information is available for this paper.

**Supplementary Figure 1:**
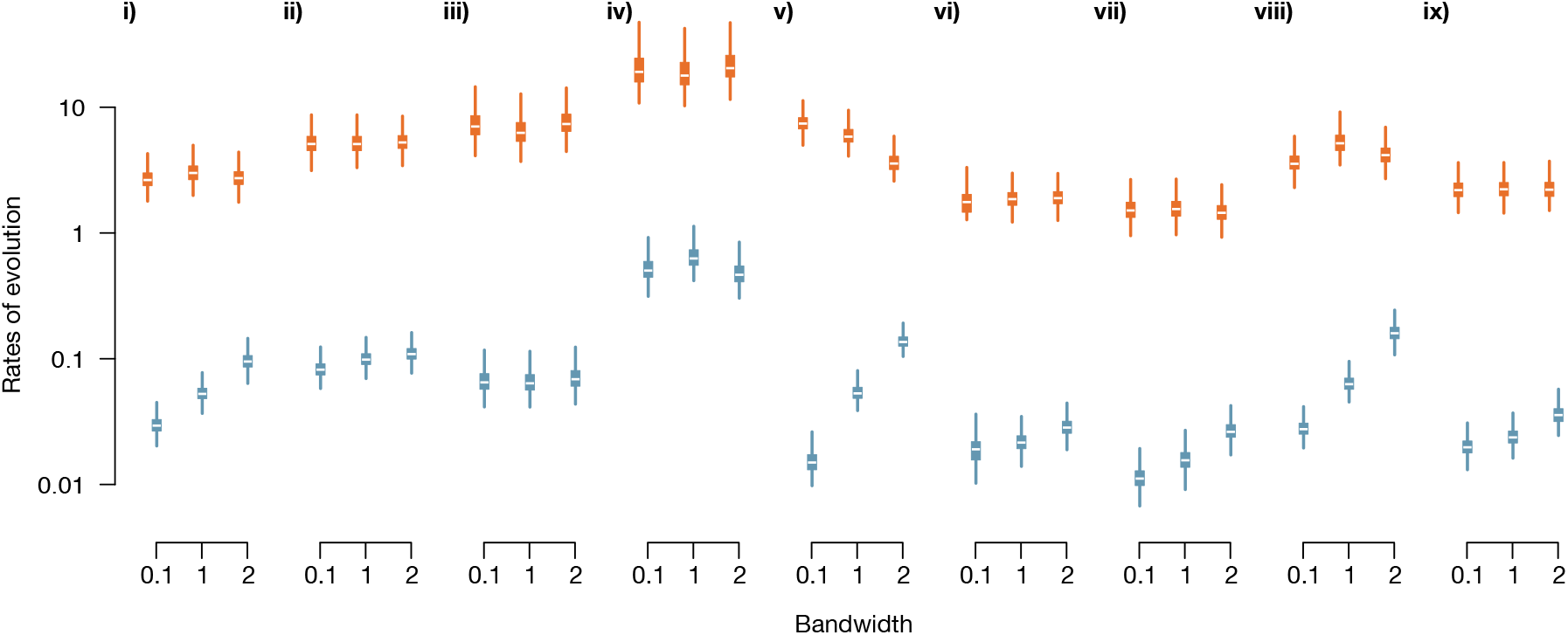
Sensitivity analyses results for the logarithm of evolutionary rates (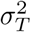 colored in orange and 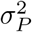 colored in light blue) using different trees (**i**-**ix**) of 18 species from the posterior and 3 different bandwidths (0.1%, 1%, 2% of the range) given by “rase”. Difference across trees display higher variation than between bandwidths, yet bandwidth selection is important to determine evolutionary rates. Overall, larger bandwidths yield higher evolutionary rates.

**Supplementary Figure 2:**
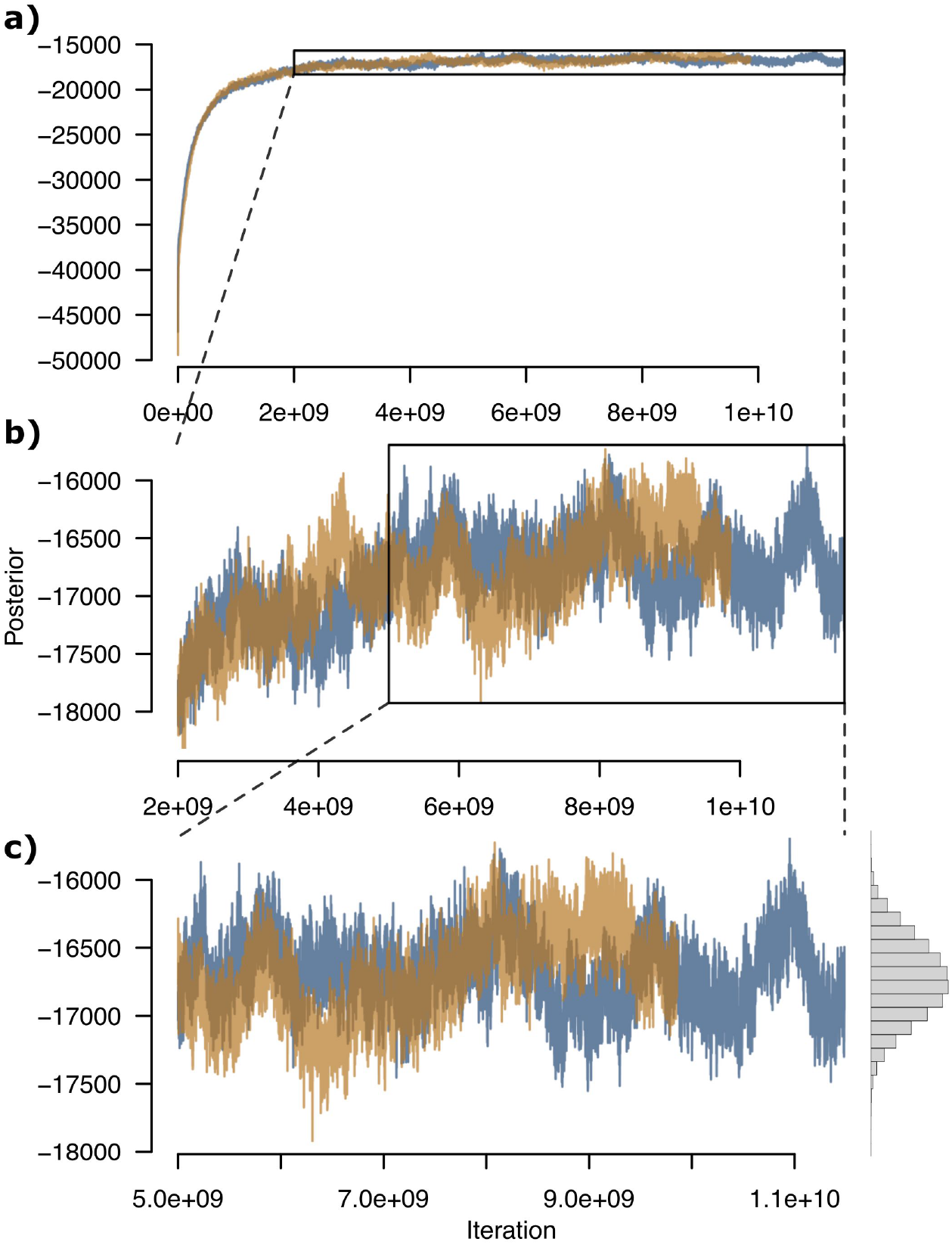
Traces of the two MCMC chains (Darwin Kernel in orange and Linux Kernel in purple) for posterior evaluations used in this paper. **a)** Trace showing the total iterations and according posterior evaluation. **b)** Trace after removing 2*e*9 iterations and according posterior evaluation. **c)** Trace after the final burn-in by removing 5*e*9 iterations, where, visually, both chains seem to have found the same stationary distribution. Histogram to the right shows the combined posterior after burn-in. Note that we do not show the thinning sampling of the final sample used for analyses.

**Supplementary Figure 3:**
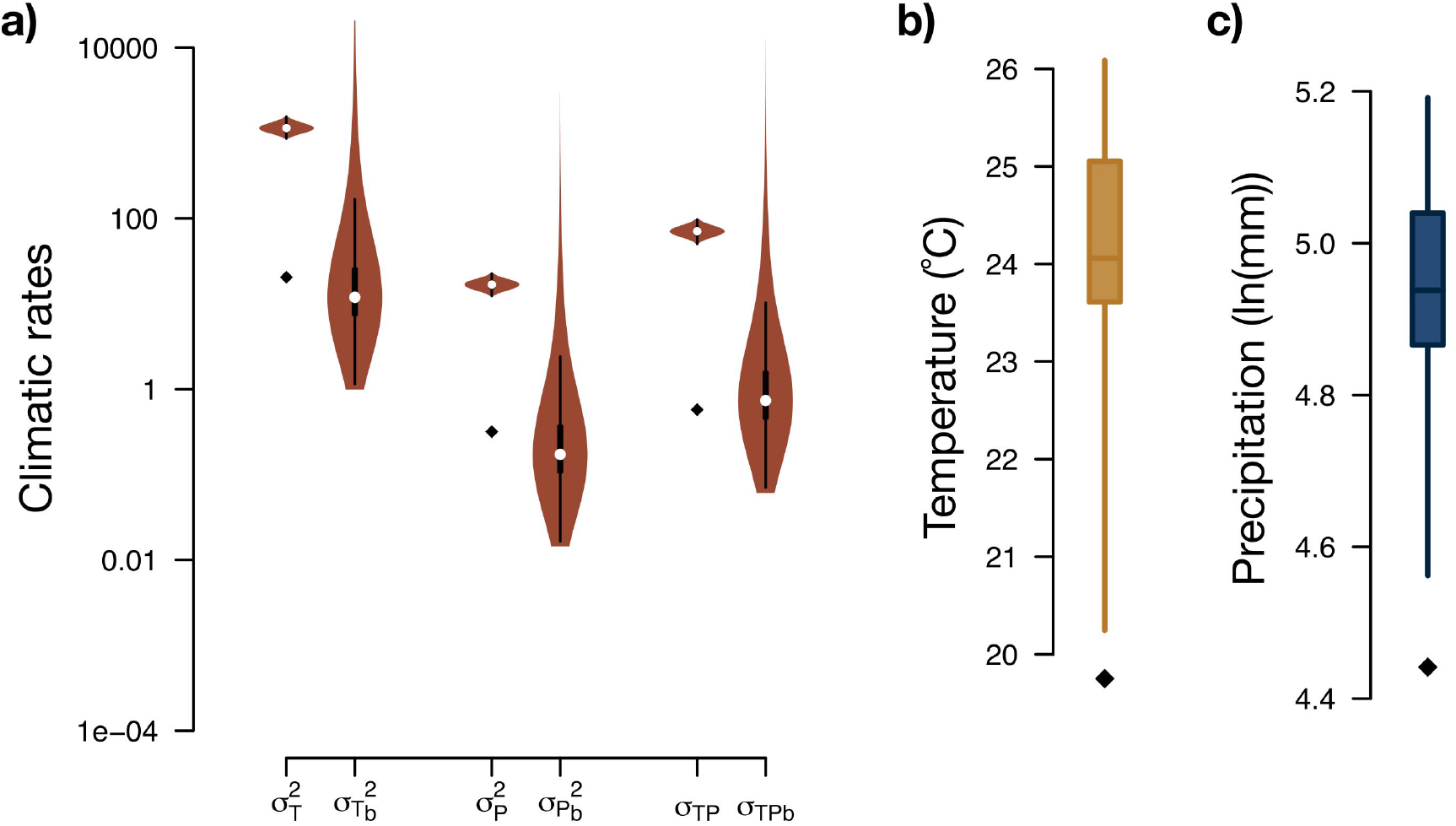
**a)** Distributions of: marginal posterior density for tree-wide rates of temperature, 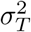, median branch-specific rates of temperature 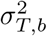, both with units ^*◦*^C^2^ My^*−*1^, marginal posterior density for tree-wide rates of precipitation, 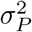, median branch-specific rates of precipitation 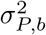, both with units ln(mm)^2^ My^*−*1^, marginal posterior density for tree-wide covariance rates 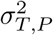 and median branch-specific covariance rates 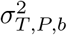. Diamonds for 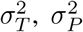 and 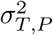 correspond to the maximum likelihood estimate (MLE) for rates of climatic evolution resulting from a multivariate strict Brownian motion (BM) using median climatic values instead of polygons (see methods). **b)** and **c)** show the marginal posterior density estimates (50% & 95% HPD) for mean annual temperature (^◦^C) and average monthly precipitation (ln(mm/month)), respectively, for the MRCA. For comparison, the diamonds correspond to the MRCA MLEs resulting from the point multivariate strict BM.

**Supplementary Figure 4:**
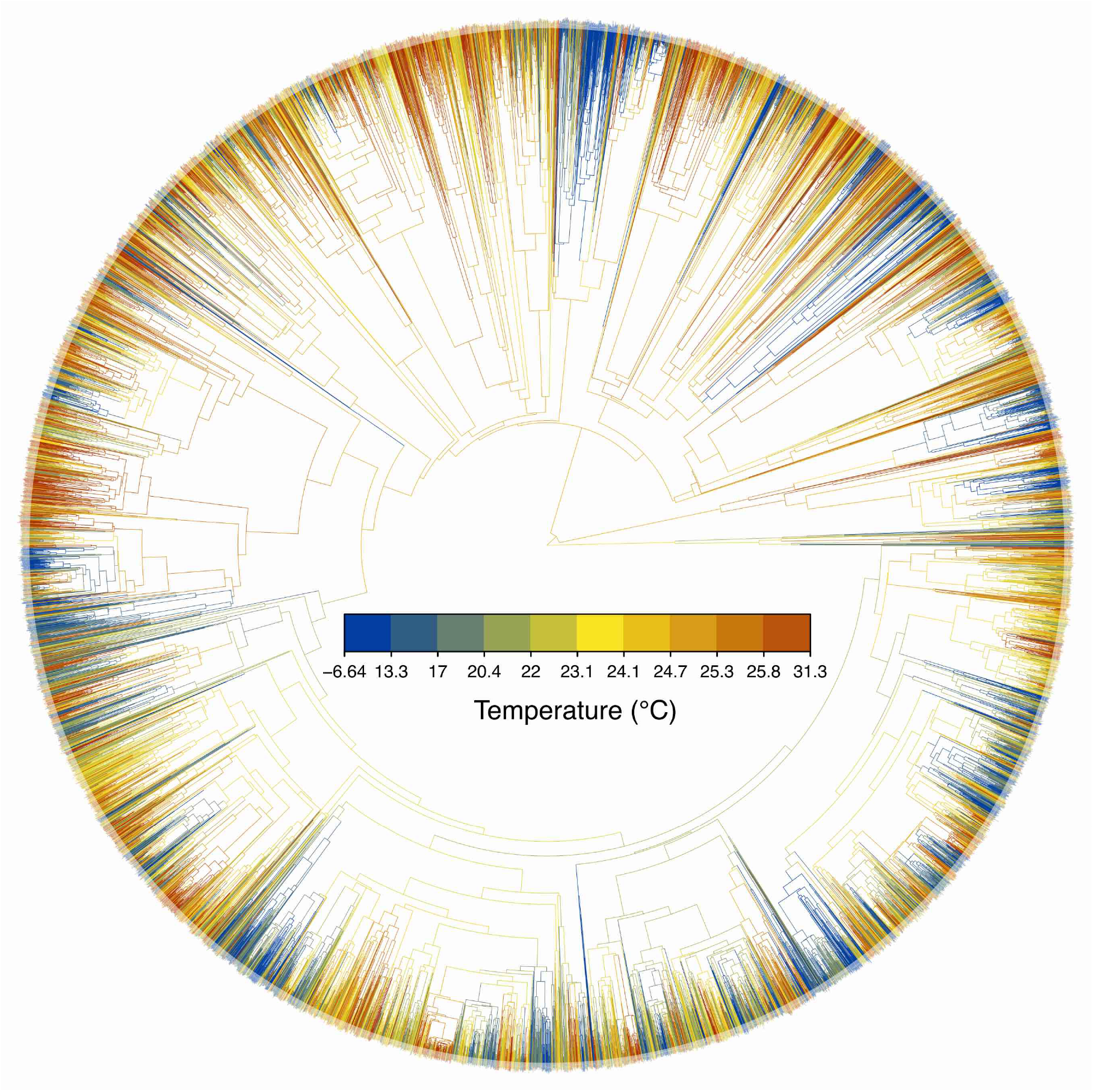
Maximum clade credibility (MCC) tree with branch coloured according to posterior median ancestral nodes estimates of average annual temperature (^*◦*^C). For visual clarity that each colour represents a posterior quantile and not equal spacing between values.

**Supplementary Figure 5:**
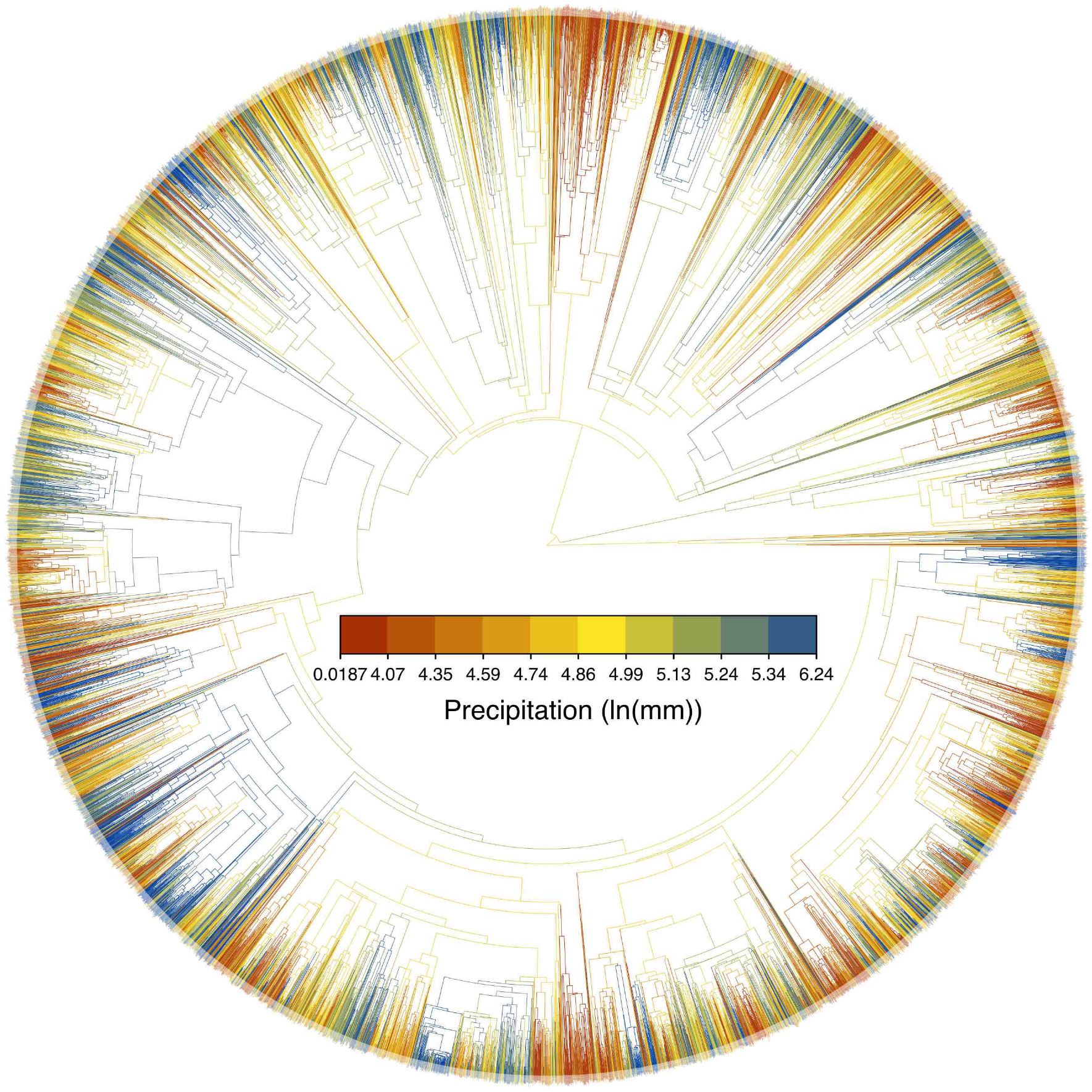
MCC tree with branch coloured according to posterior median ancestral nodes estimates average monthly precipitation (ln(mm)). For visual clarity that each colour represents a posterior quantile and not equal spacing between values.

**Supplementary Figure 6:**
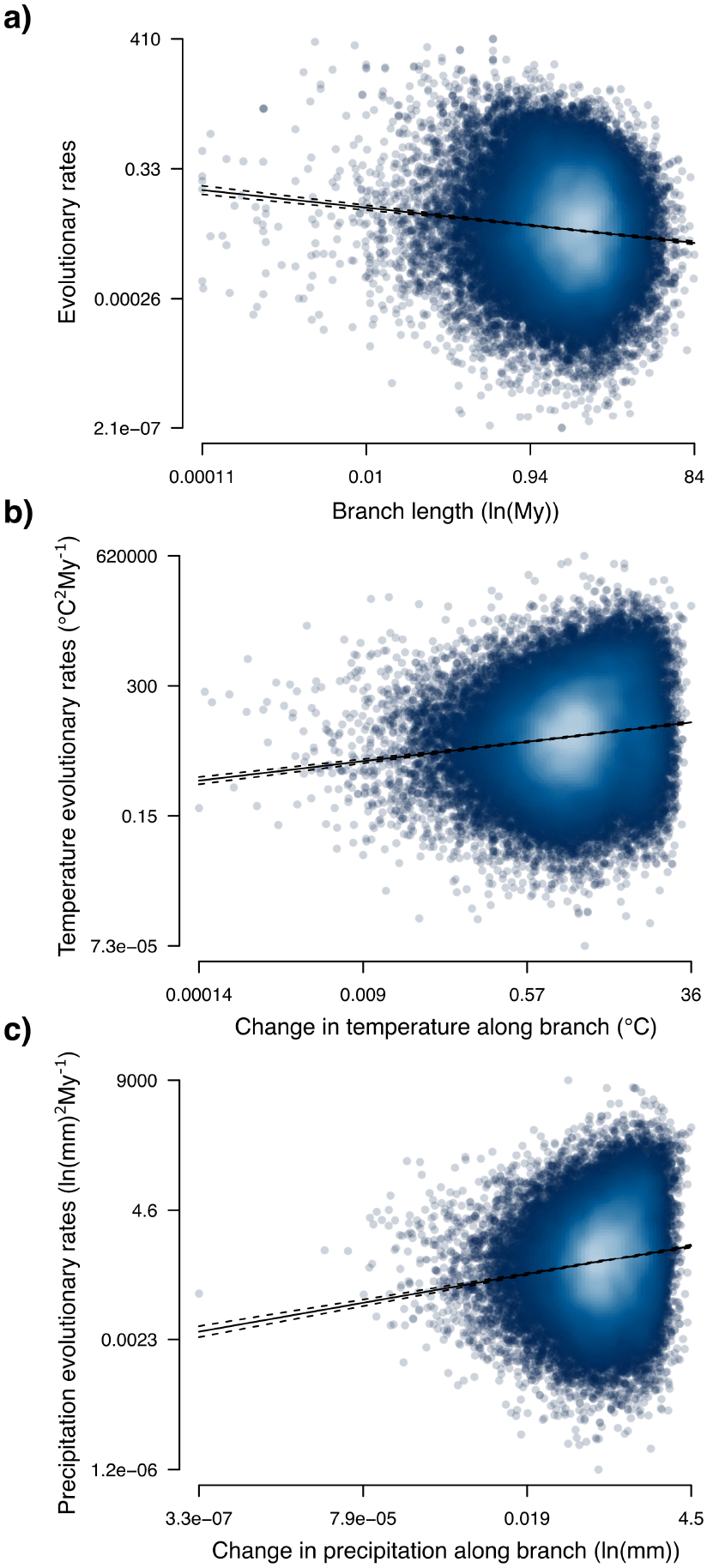
**a)** Relationship between the branch-specific posterior median of rates of climatic niche evolution, *ϕ*_*b*_, and branch length both transformed to natural logarithms. **b)** Relationship between the branch-specific posterior median of rates of temperature niche evolution, *ϕ*_*b*_, multiplied by branch length and measured change in temperature along the branch. **c)** Relationship between the branch-specific posterior median of rates of precipitation niche evolution, *ϕ*_*b*_, multiplied by branch length and measured change in precipitation along the branch. In all plots darker tones represent higher posterior density. Black solid line is the best fit linear model and dashed lines denote the 95% confidence interval.

**Supplementary Figure 7:**
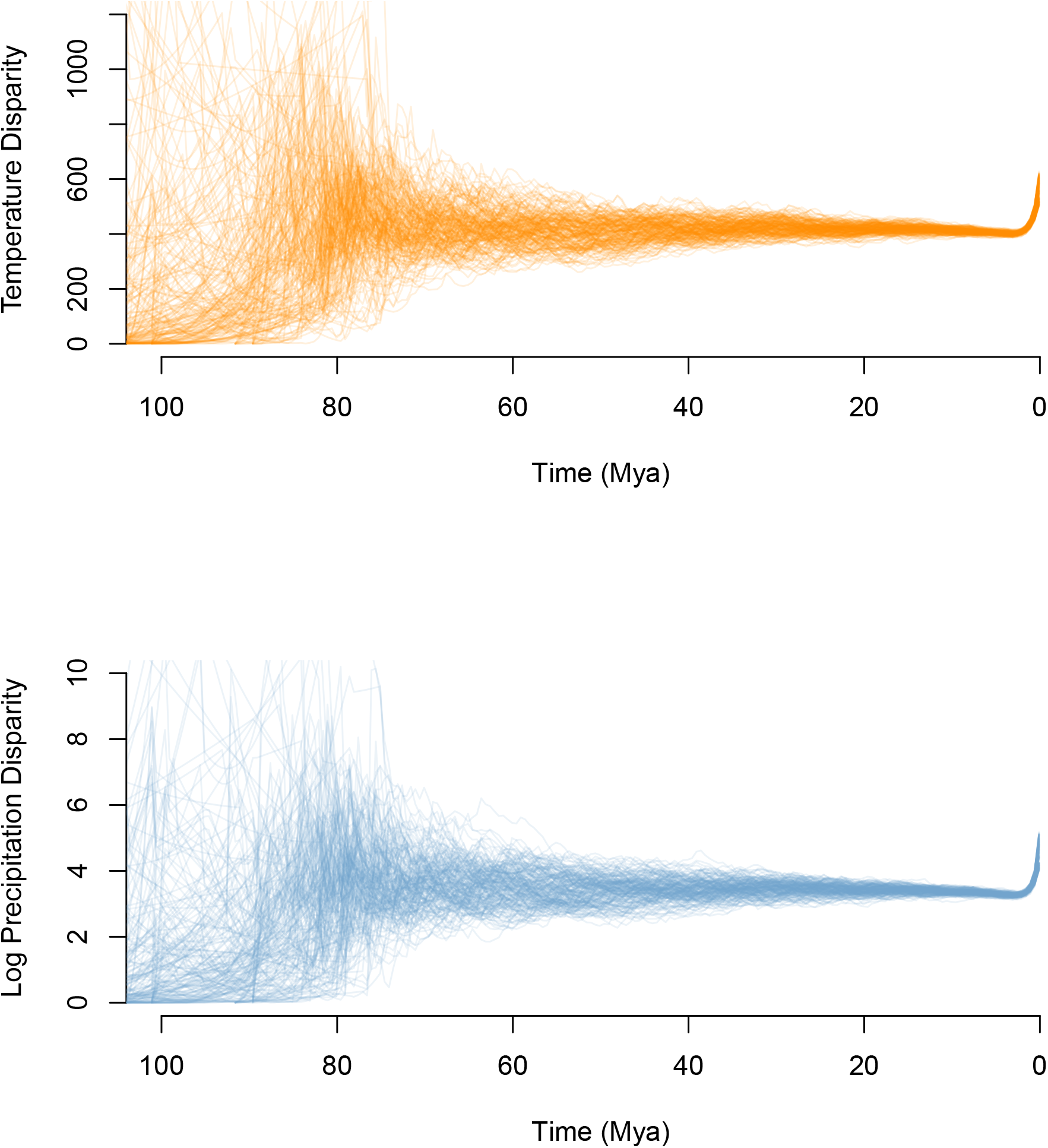
10 Bounded evolution simulations of disparity on each the 100 trees for **top)** Temperature, and, **bottom)** Log Precipitation.

## Supplementary Videos

**Video S1:** Environmental space showing niche filling throughout the Aves radiation for a randomly sampled tree. The hexagonal grid is drawn by summarizing each lineage’s climatic niche as a 95% highest posterior density (HPD) polygon and coloring each grid cell by with the count of overlapping climatic polygons at a given time. As a point of reference, the currently occupied realized climatic space for all birds is shown in grey background.

**Video S2:** Environmental space showing climatic niche evolutionary rates throughout the Aves radiation for a randomly sampled tree. The hexagonal grid is drawn by summarizing each lineage’s climatic niche as a 95% highest posterior density (HPD) polygon and coloring each grid cell by the median evolutionary rate of overlapping climatic polygons at a given time. As a point of reference, the currently occupied realized climatic space for all birds is shown in grey background.

## Notes

### Competing Interest Statement

The authors have declared no competing interest.

